# G3DCT: An Interpretable Spatial Grid-based Framework with Temporal Convolution-Transformer for EEG Artifact Identification

**DOI:** 10.64898/2026.01.30.702940

**Authors:** Aonan He, Xi Wang, Jiangwei Yu, Xiaojia Wang, Zongyuan Ge, Youyong Kong, Guanyu Yang, Chunfeng Yang, Chen Yang, Miao Cao

**Affiliations:** School of Computer Science and Engineering, Southeast University, Nanjing, 210096, Jiangsu, China; School of Biological Science and Medical Engineering, Southeast University, Nanjing, 210096, Jiangsu, China; School of Health Sciences, Swinburne University of Technology, John Street, Hawthorn, Victoria, 3122, Australia; Department of Electrical and Computer Systems Engineering, Monash University, Clayton, Victoria, Australia; Wuxi Vocational College of Science and Technology, Wuxi, 214028, China

**Keywords:** Electroencephalography, Artifact Identification, Deep Learning, Attention Mechanism, Focal Loss

## Abstract

Electroencephalography (EEG) serves as a fundamental tool in modern neurology, cognitive neuroscience, and brain-computer interfaces, but its practical application is often compromised by artifacts. Physiological artifacts are particularly intractable due to overlapping spectral features with neural signals, hindering reliable EEG interpretation. In this work, we propose Grid-based 3D Convolution-Transformer (G3DCT), an interpretable deep learning framework for EEG artifact identification. The framework embeds multi-channel EEG signals into fixed grids to leverage electrode spatial topology, employs parallel multi-branch temporal convolutions and Transformers to handle complex artifacts, and incorporates an attention module to visualize scalp activation patterns, which enhances physiological interpretability. Our evaluation on three datasets demonstrates that G3DCT outperforms existing state-of-the-art models. For challenging combined artifacts, it secures a gain of 2.8% in F1-score over the second-best model. These results demonstrate that G3DCT provides an efficient and robust solution for EEG artifact identification, which has the potential to enhance the reliability of EEG-based applications in practice.

## 1. Introduction

Electroencephalography (EEG) is fundamental to neurology 1, neuro-science 2, and applications such as brain–computer interfaces (BCIs) 3. However, EEG signals are more susceptible to artifacts, compared with other functional imaging modalities including magnetoencephalography and functional magnetic resonance imaging. These artifacts can originate from instrumental sources (e.g., electrode failures, power-line interference) or physiological sources (e.g., eye movements, blinks, muscle activity) 4. Instrumental artifacts can often be reduced by recording in a controlled environment and applying simple filtering, whereas physiological artifacts are difficult to distinguish and remove because their spectral content frequently overlaps the desired signal and they show considerable variation across different origins 5. Such artifacts can obscure genuine neural activity 5, interfere with the decoding of neural information, and ultimately lead to misinterpretations of EEG data 6. Notably, a comprehensive review covering 90 related studies revealed that 28% relied on manual artifact removal, while another 22% did not apply any correction procedures 7. This highlights the current lack of unified and standardized artifact processing protocols in the field.

Traditional signal processing techniques, such as regression analysis [8, 9], blind source separation (BSS) [10, 11], and frequency-domain filtering [12, 13], remain valuable for their strong interpretability. However, these approaches often rely on prior information and manual input, such as the need for high-quality reference channels 14, manual selection of independent components 15, or setting filtering thresholds 16, making it difficult to meet the demands of real-time processing and automation. In recent years, researchers have explored hybrid strategies, such as empirical mode decomposition (EMD)–independent component analysis (ICA) 17, Wavelet–ICA 18, and BSS–support vector machine (SVM) 19, to improve the accuracy and automation of artifact identification. Nevertheless, due to their high computational cost, these methods are primarily suited for offline scenarios 4.

Prior to the rise of deep learning, traditional machine learning methods combined with manual feature extraction formed a fundamental basis for EEG artifact recognition. These approaches typically involved extracting features such as frequency band energy and inter-channel correlation matrices, which were then used to train conventional classifiers like Linear Discriminant Analysis (LDA) 20 and k-Nearest Neighbors (k-NN) 21. However, their ability to generalize across subjects and datasets is often limited, as inter-subject variability and differences in artifact induction protocols can shift the temporal and spectral characteristics of artifacts. More critically, manual feature extraction is constrained by the researcher’s prior knowledge, introducing subjectivity and making it difficult to capture the underlying feature patterns of artifacts. For example, Roy et al. 22 reported that multiple classifiers failed to adequately recognize shivering artifacts when provided with spectral features derived from the Fast Fourier Transform (FFT).

In contrast, deep learning demonstrates the capability to learn effective features directly from EEG data without hand crafting. Khatwani et al. 23 designed a lightweight CNN architecture that enables efficient artifact detection by processing two-dimensional EEG representations. To enhance cross-device generalization performance, van Stigt et al. 24 employed transfer learning to fine-tune a CNN model on dry-electrode EEG data, achieving an artifact recognition accuracy of 90.7%. To capture long-range dependencies, Semkiv et al. 25 built a hybrid CNN–LSTM model with an attention mechanism in a single-channel EEG study, improving the precision of artifact detection and localization. Gao et al. 26 further proposed a dual-scale CNN–LSTM model to reconstruct clean EEG signals. With the growing need for global context modeling, the Transformer architecture has also gained attention in EEG artifact recognition. Chuang et al. 27 proposed a Transformer-based model for simultaneous multi-class artifact recognition and reconstruction, while Tang et al. 28 introduced a CNN–Transformer dual-stage network (CT-DCENet) to advance EEG denoising performance.

Previous studies have systematically explored various deep learning architectures, ranging from local feature extraction with CNNs to long-range dependency modeling. However, the adaptability and robustness of current models in real-world scenarios still face a series of challenges. Firstly, spatial layout information of electrodes is often underutilized. Most studies 7, 24, 27, 29, 30 employ input representations such as ‘channels × time points’ matrices or single-channel time series, which fail to capture the spatial topological relationships between electrodes. Secondly, research on modeling combined artifacts remains limited 31. In real-world recordings, artifacts frequently co-occur rather than appearing in isolation 6. For example, head movements may coincide with eyebrow raising. However, existing work 32–34 has primarily focused on isolated artifact types like eye blinks or muscle activity. Thirdly, model interpretability is often inadequate 35, 36. Many deep learning approaches 37, 38 operate as ‘black-box’ systems, providing limited physiologically grounded explanations for their predictions.

In this study, we addressed the aforementioned challenges by introducing a unified deep learning framework with the following key designs: (1) standardizing the input by embedding multi-channel EEG into a fixed grid to leverage electrode topology; (2) constructing a hybrid architecture that combined temporal convolutional pyramids (TCP) and pre-normalized Transformers to effectively model complex combined artifacts; and (3) incorporating a Fast Scalp Attention Module (FSAM) to visualize scalp activation regions and provide physiological explanations for model decisions. The overall framework is illustrated in Fig. 1.

**Figure 1.**
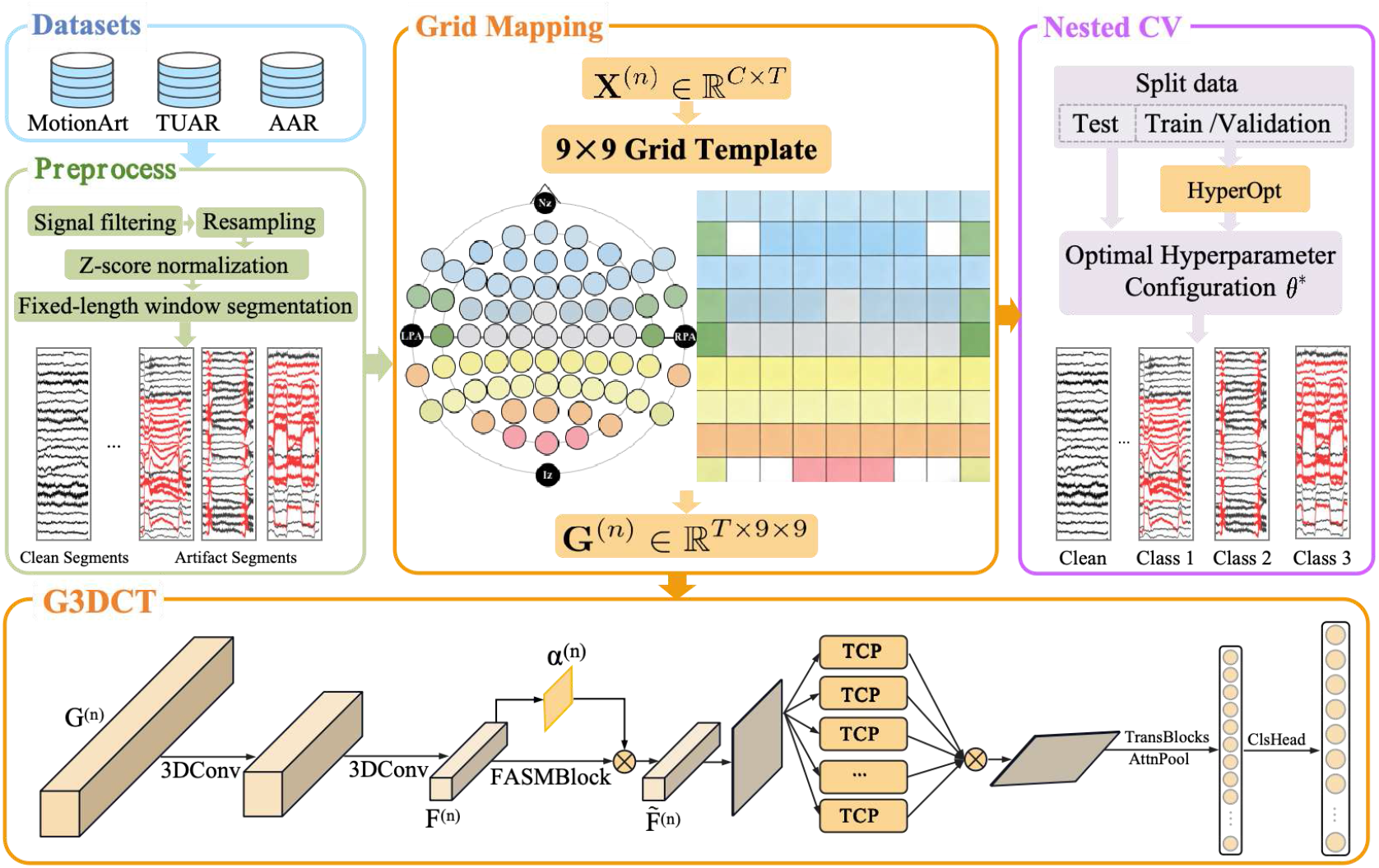
Overview of the G3DCT framework for EEG artifact identification. The processing pipeline begins with the preprocessing of three public datasets (MotionArt, TUAR, and AAR), followed by the projection of EEG signals onto a fixed grid based on electrode positions. The lower subfigure details the sequential components of the model architecture, including three-dimensional convolution (3DConv), Fast Spatial Attention Module (FSAM), Temporal Convolutional Pyramid (TCP), Transformer blocks (TransBlocks), attention pooling (AttnPool), and classification head (ClsHead). Model evaluation employed nested cross-validation (nested CV) with grouped *K*-fold cross-validation (GroupKFold) to prevent data leakage, while hyperparameters were optimized using Bayesian hyperparameter optimization (HyperOpt).

The primary contributions of this work are summarized as follows:

- We proposed Grid-based 3D Convolution-Transformer (G3DCT), an interpretable deep learning framework that incorporates electrode spatial topology for EEG artifact identification.
- We evaluated the proposed model on three public datasets. The experimental results demonstrate that our method outperforms existing state-of-the-art models in overall performance.
- We showed that the scalp regions localized by the FSAM align with the known physiological origins of artifacts, enhancing physiological interpretability.
- We demonstrated that explicitly modeling the spatial relationships among electrodes improves the discriminability between different artifact types.

## 2. Methods

### 2.1. Data Representation and Topographic Embedding

Each artifact exhibits a characteristic spatial pattern, and even artifacts with similar waveforms can show different distributions across the scalp. Fig. 2 illustrates the temporal waveforms and corresponding scalp topographies of various artifact types. This highlights the importance of effectively modeling inter-electrode spatial relationships for accurate identification. However, variations in EEG montages and recording devices make it difficult to leverage such spatial information consistently. To overcome this limitation and better exploit the inherent electrode topology, we embedded signals from diverse datasets into a unified grid template. During preprocessing, continuous EEG signals were segmented with a fixed window length *T* and a given stride. The *n*-th segment was denoted as **X**^(n)^ ∈ ℝ^*C×T*^, where *C* was the number of EEG channels. As shown in Fig. 3, each channel was uniquely mapped to a coordinate on a 9 *×* 9 grid according to its anatomical location, with missing positions filled by zero-padding. This procedure converted each segment into a standardized 3D tensor **G**^(n)^ ∈ ℝ^*T ×*9*×*9^, which subsequently served as the model input.

**Figure 2.**
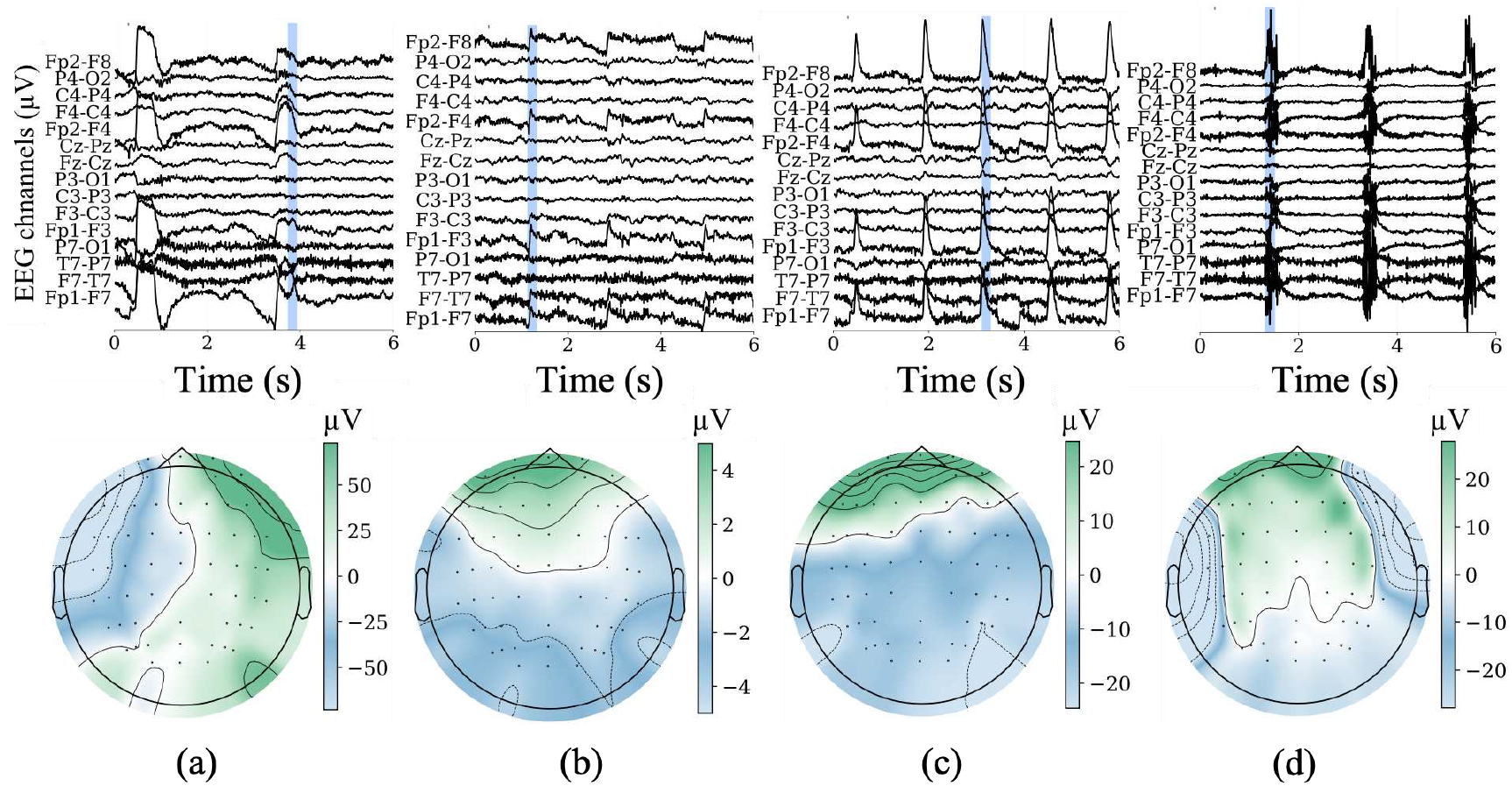
Temporal waveforms and scalp topographies of representative EEG artifacts. The blue-shaded interval (0.20 s) in the waveforms denotes the key period of artifact occurrence, with the corresponding scalp potential distribution shown below. (a) Horizontal head movement; (b) Vertical eye movement; (c) Eye blink; (d) Chew.

**Figure 3.**
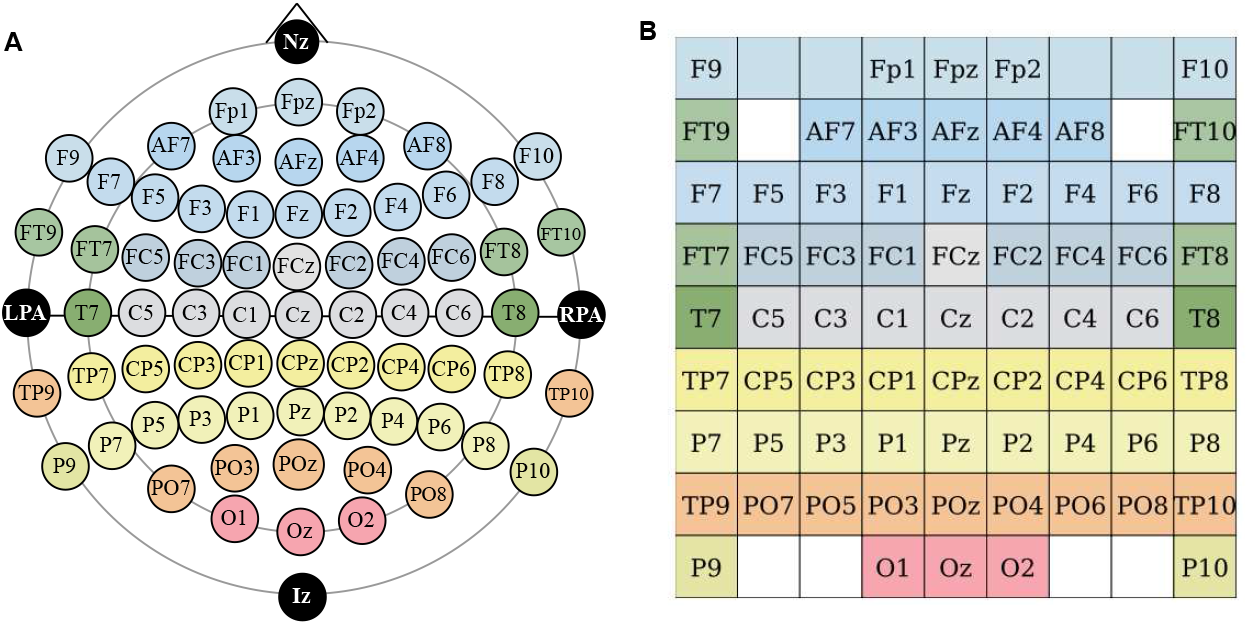
Standardized EEG representation via grid mapping. (A) Electrode locations follow the international 10–10 system. (B) Channels from different datasets are mapped onto a common 9 *×* 9 grid template, which incorporates 71 commonly used channels. MotionArt contains 59 channels, all of which are included in the template. The AAR dataset consists of 30 channels, 27 of which are present in the template, excluding EOG1, EOG2, and EOG3. For the TUAR dataset, which involves more complex channel configurations, we selected the 19 common channels present in all its recordings, each of which is mapped to the template. Detailed channel information for each dataset is provided in the Supplementary Material.

### 2.2. Model Architecture

#### 2.2.1. Local Spatiotemporal Feature Extraction

The model was designed to process the standardized grid representation **G**^(n)^ and hierarchically extract features for artifact identification. Artifacts in EEG signals frequently manifest as localized scalp patterns that evolve dynamically over brief intervals. To effectively capture these distinctive spatiotemporal signatures, we employed a residual 3D Convolutional Neural Network (CNN), as shown in Fig. 4B. The architecture consisted of two residual blocks. The first block primarily reduced the temporal length while preserv-ing the full spatial grid, and the second block further compressed both the temporal and spatial dimensions. Residual connections were incorporated to facilitate gradient flow and stabilize the training of deeper layers. The input tensor **G**^(*n*)^ was processed by this network to produce a 4D feature tensor 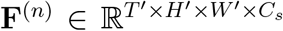, where *T* ^*′*^ was the downsampled temporal length, *H*^*′*^ and *W* ^*′*^ were the reduced spatial dimensions, and *C*_*s*_ was the number of feature channels.

**Figure 4.**
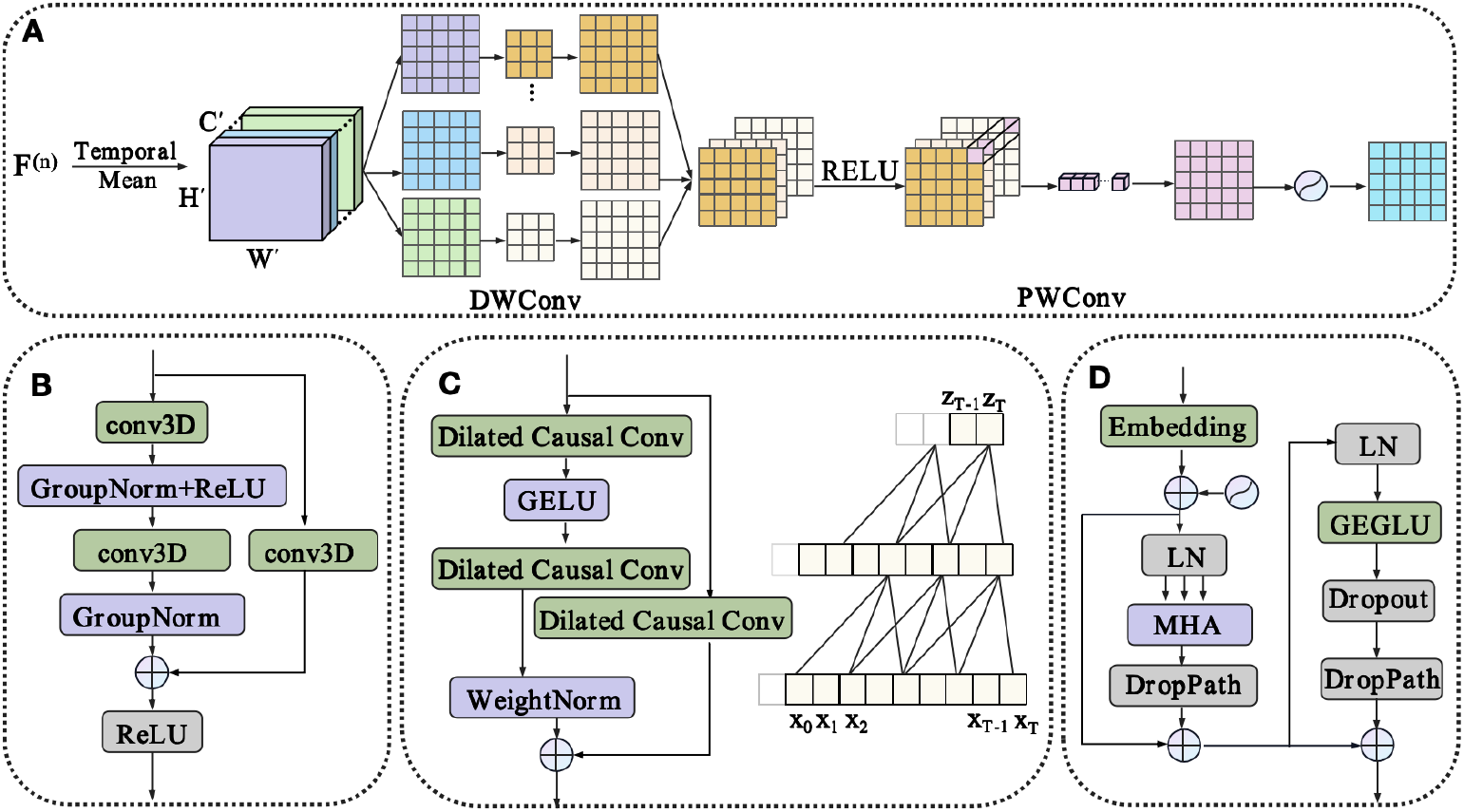
Overview of the main blocks of G3DCT. (A)FSAM generates spatial attention by first aggregating temporal information and then applying separate depthwise and pointwise convolutions to identify salient scalp regions. (B) Residual 3D CNN extracts local spatiotemporal features. (C) TCP captures multi-scale temporal dependencies using parallel dilated temporal convolutions with kernel sizes 3,7,15 and dilation rates 1,2,4. (D) Transformer block performs global temporal modeling via a pre-normalized Transformer equipped with multi-head attention. DWConv: depthwise convolution; PWConv: pointwise convolution; conv3D: 3D convolution; ReLU: rectified linear unit; GroupNorm: group normalization; GELU: Gaussian error linear unit; WeightNorm: weight normalization; LN: layer normalization; MHA: multi-head self-attention; GEGLU: gated GELU; DropPath: stochastic depth (drop-path); Embedding: embedding layer.

To direct the model’s focus to scalp regions that are physiologically relevant to artifact generation, we introduced the Fast Scalp Attention Module (FSAM), illustrated in Fig. 4A. This module provided a learnable mechanism to emphasize critical spatial areas. Specifically, temporal information was first aggregated by averaging over time:

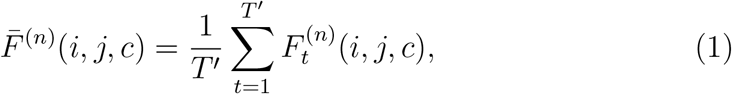

where 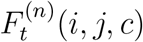 was the feature value at time *t*, grid position (*i, j*), and channel *c*. The aggregated feature map 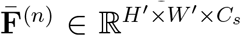 was then passed through a shallow convolutional block followed by a sigmoid activation, generating a spatial attention weight map *α*^(n)^(*i, j*) ∈ (0, 1). This weight map was broadcast to all time steps to recalibrate the original features:

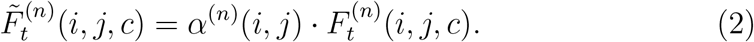

Consequently, features from scalp regions more indicative of artifacts received higher weighting for subsequent processing. To prepare for the subsequent temporal modeling, the spatially weighted features at each time step were flattened and linearly projected to a common dimensionality *d*. This was achieved by a learned linear projection:

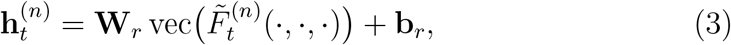

where vec(·) flattened the (*H*^*′*^, *W* ^*′*^, *C*_*s*_) dimensions, and 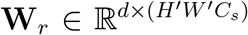,**b**_*r*_ ∈ ℝ^*d*^ were learnable parameters. Stacking the projections from all time steps yielded the sequence:

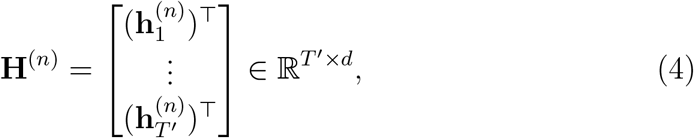

which served as the input to the subsequent global temporal modeling stage.

#### 2.2.2. Multi-scale Temporal Modeling

Artifacts exhibit dynamic patterns that operate across multiple temporal scales, ranging from brief muscular contractions to sustained movements. To effectively model this hierarchical temporal structure, we first processed the feature sequence **H**^(n)^ with a Temporal Convolutional Pyramid (TCP) module. This module applied multiple parallel 1D convolutional branches with varying kernel sizes and dilation rates to the input sequence (Fig. 4C):

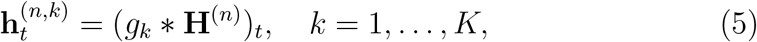

where *g*_*k*_ denoted the convolutional kernel of the *k*-th branch. The outputs were combined and projected to produce the enhanced sequence 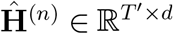

While TCP modules are effective at capturing local temporal patterns, their finite receptive field limits their ability to model long-range dependencies within the sequence, often necessitating deep architectures for long-range dependencies 39, 40. To overcome this limitation and capture global dependencies across the entire time series, we employed a Transformer encoder 41. To preserve the sequential order of features, a learnable positional encoding **P** ∈ ℝ^T ′ × d^was first added to the input (Fig. 4D):

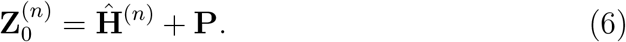

The sequence was then processed by *L* pre-normalized Transformer blocks. Each block updated the features as:

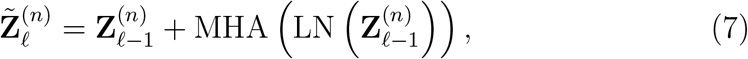

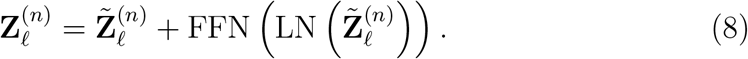

where MHA denoted multi-head self-attention, FFN was a feed-forward network, and LN was layer normalization. The final output 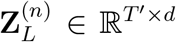 in-tegrated both local multi-scale features and global context, with 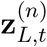 representing the feature at time step *t*.

#### 2.2.3. Attention Pooling and Classification Head

To aggregate the temporal features into a single vector representation for the entire segment, we employed an attention-based pooling mechanism. This allowed the model to dynamically emphasize the most salient time steps for the final classification. Specifically, a learnable query vector **q** ∈ ℝ^*d*^ was used to compute an attention score for each time step:

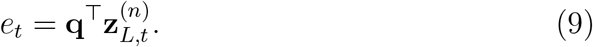

The scores were normalized across the temporal dimension via a softmax function to obtain the attention weights:

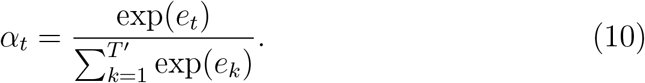

The segment-level representation was then computed as a weighted sum of the temporal features:

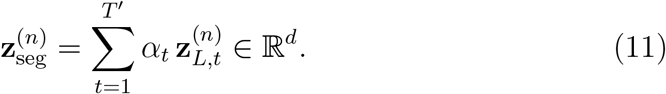

Finally, this representation was passed through a multilayer perceptron classification head to produce the predicted probability distribution over the *K* artifact classes:

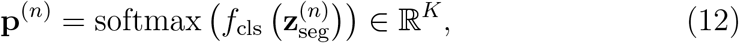

where *f*_cls_( ) was a feed-forward network consisting of multiple fully connected layers.

### 2.3 Training Strategy and Hyperparameter Optimization

The annotation of EEG artifacts often results in a long-tailed class distribution, where certain artifact types occur far less frequently than others. To mitigate the bias introduced by such class imbalance during training, we adopted Focal Loss as the optimization objective. Compared with other loss functions for class imbalance mitigation (e.g., weighted cross-entropy), Focal Loss reduces the contribution of well-classified examples in an adaptive way, thereby focusing optimization on difficult and misclassified instances and reducing sensitivity to manually tuned static class weights 42. For a single sample (**X**^(n)^, *y*^(n)^), the Focal Loss was defined as:

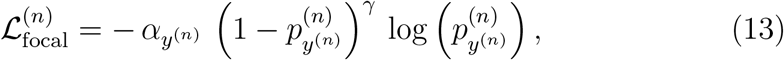

where 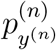 denoted the predicted probability for the ground-truth class *y*^(n)^,*γ >* 0 was the focusing parameter that controls the down-weighting of easy examples, and was a class-specific weighting factor that counteracts frequency imbalance. The class weight *α*_*k*_ for class *k* was constructed in two steps. First, an inverse-frequency weight was computed:

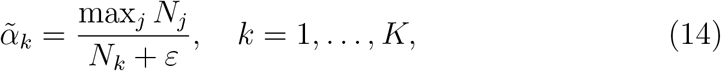

where *N*_*k*_ denoted the number of training samples in class *k*, max_*j*_ *N*_*j*_ represented the sample count of the largest class, and *ε* was a small constant (e.g., 10^−8^) added for numerical stability. These base weights were then adjusted by a learnable scaling factor *s*_*k*_ and a global scaling coefficient *λ*. The final normalized class weight was:

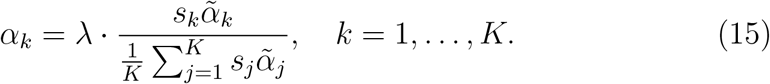

To ensure an unbiased evaluation during hyperparameter optimization, we adopted a nested cross-validation scheme. The procedure consisted of an outer loop and an inner loop. In the outer loop, the dataset was partitioned into *K*_outer_ folds and each fold served once as the test set while the remaining folds were used for model development. Within each outer fold, the inner loop performed hyperparameter search using Tree-structured Parzen Estimators (TPE). The TPE algorithm minimized the inner-loop objective:

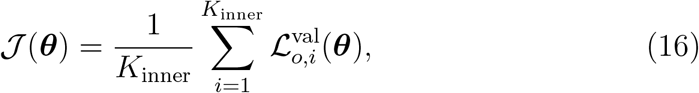

where 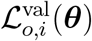 denoted the validation loss obtained with hyperparameters ***θ*** on the *i*-th inner validation split of the *o*-th outer training fold. The hyperparameter set ***θ***^∗^ that minimized 𝒥 (***θ***) was selected to train the final model for evaluation on the outer test fold.

## 3 Experiments

### 3.1 Datasets

To comprehensively assess the generalizability and robustness of the proposed method, we employed three complementary EEG artifact datasets. These datasets exhibit significant differences in channel counts, artifact categories, recording environments, and degrees of class imbalance. Detailed descriptions of recording protocols and label distributions are provided in the Supplementary Material.

**MotionArt** is a dedicated dataset constructed for this study. During data collection, participants performed predefined actions such as eye blinks, tongue movements, and head rotations. Channel-wise labels were annotated by experts into 14 classes, including both individual artifact types and their combinations. In addition, the dataset also includes clean EEG segments, enabling its use for both artifact detection and classification.

**TUAR** 43 represents a large-scale and real-world clinical environment, containing diverse common artifacts like muscle activity (Musc), eye movements (Eyem), and combined categories. Its highly imbalanced data distribution poses a significant challenge for model generalization and robust evaluation.

**AAR** 44 was also acquired under controlled laboratory conditions. It contains nine artifact types with a relatively balanced class distribution.

### 3.2 Preprocessing and Segmentation

A standardized preprocessing pipeline was applied to all raw EEG recordings. Signals were first notch-filtered at 50 or 60 Hz to remove line noise and then high-pass filtered at 1 Hz to eliminate slow drifts. All data were downsampled to a common sampling rate of 125 Hz. For temporal segmentation, a fixed window length of *T* = 3 s with a stride of 1.5 s was employed. To increase the usefulness of training samples, we designed a selective segmentation strategy. Let an annotated artifact interval be defined as

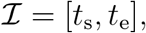

where *t*_s_ and *t*_e_ denote its start and end times. To ensure that brief artifacts such as vertical eye movements are not missed or excessively truncated by the sliding windows, the first window was anchored at the onset of the interval:

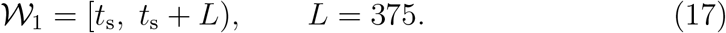

Subsequent windows were then generated by advancing in steps of half the window length:

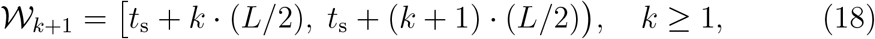

until *t*_s_ +(*k* +1) · (*L/*2) *> t*_e_. This procedure guaranteed that each labeled artifact interval contributed at least one representative segment, while avoiding excessive inclusion of non-artifactual signal. Since sliding-window segmentation produces overlapping data segments, potential leakage can arise from the high similarity of adjacent windows. To prevent this, we ensured that data from any single subject never appeared simultaneously in the training and test sets.

### 3.3 Baseline Methods

To provide an objective evaluation of the proposed G3DCT model, its performance was compared against several representative baseline methods. The selected models are widely used in EEG classification and have achieved competitive results in prior studies. To ensure a fair comparison, all baselines followed the same preprocessing and data partitioning protocol. The optimizer and loss function were kept identical across all models. Variations were permitted only in the core network architecture and its corresponding parameters. Complete implementation details for each baseline are provided in the Supplementary Material.

**CLAttn** (CNN–LSTM with Attention) 45 was proposed by Cisotto et al. for EEG artifact detection on the Temple University Hospital (TUH) dataset. The model integrates a convolutional neural network (CNN) for feature extraction, a Long Short-Term Memory (LSTM) network for temporal modeling, and an attention mechanism to emphasize informative time features, achieving strong performance in prior studies.

**TCN** (Temporal Convolutional Network) 40 was introduced by Bai et al., demonstrating superior performance and stability over traditional recurrent neural networks (RNNs) on various sequence modeling baselines. Its application was extended to EEG analysis by Gemein et al. 46, who utilized TCNs to attain high accuracy in detecting pathology within the TUH abnormal EEG corpus.

**GAT** (Graph Attention Network) 47 employed attention mechanisms to adaptively weight the importance of connections between nodes in a graph. Demir et al. 48 applied this architecture to EEG classification by using spatial features from specific frequency bands, showing that their EEG-GAT model outperformed other contemporary graph neural network models.

**RNN-CNN** 49, 50 hybrid approaches are common in EEG analysis. A review by Roy et al. 49 noted that from 2010 to 2018, approximately 40% of deep learning EEG studies used CNNs and 14% used RNNs. A notable implementation by Kim et al. 50 integrated a recurrent branch with multiple 1D convolutional branches, achieving 67.6% accuracy on a five-class artifact task in the TUH dataset while maintaining a very fast inference time.

**LDA** (Linear Discriminant Analysis) 51 represents a strong traditional machine learning baseline methods for EEG artifact classification. Roy 22 reported that LDA achieved the best artifact recognition performance on the TUH dataset compared to other classical methods when combined with hand-crafted features.

### 3.4 Implementation Details

All experiments were conducted on a Linux server using Python and PyTorch on NVIDIA GPUs. To enhance training efficiency, mixed-precision training was employed and a subset of the G3DCT modules were frozen during the initial epochs. In the inner loop of nested cross-validation, key hyperparameters including batch size, initial learning rate, weight-decay coefficient, the focal loss focusing parameter *γ*, and the per-class scaling factors {*s*_*k*_} were tuned automatically. The search ranges for these hyperparameters and the optimal configurations are detailed in the Supplementary Material.Training for any candidate configuration was stopped if the validation loss did not improve for five consecutive epochs with a maximum of 50 epochs per fold. Once the optimal hyperparameters 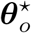 were selected in the inner loop, the model was retrained in the outer stage using the combined training and validation sets. To better exploit the available data, early-stopping patience was relaxed to eight epochs in this phase. Across all methods, the AdamW optimizer with weight decay was used to mitigate overfitting.

## 4. Results

We first evaluated the proposed Grid-based 3D Convolution-Transformer (G3DCT) model against representative baseline methods by performing artifact detection on MotionArt and artifact classification across all three datasets. Next, the robustness of G3DCT was examined through a sensitivity analysis of four key hyperparameters. Finally, an ablation study was conducted to quantify the contribution of each major component within the G3DCT framework. The key findings are summarized in the following sections, while extended results and supporting analyses are provided in the Supplementary Material. Evaluation metrics included accuracy (ACC), precision (Pre), recall (Rec), F1-score (F1), specificity (Spe), balanced accuracy (BAC), and weighted F1-score.

### 4.1. Artifact Classification Performance

On the MotionArt detection task, G3DCT achieved an accuracy of 97.3%, surpassing all baseline models. Detailed results are reported in Table 1. In the more challenging multi-class artifact classification setting, G3DCT consistently delivered superior performance across all three datasets (Table 2). Notably, it achieved the best scores on all metrics for MotionArt and TUAR, both datasets characterized by complex combined artifacts and imbalanced class distributions. On MotionArt, G3DCT attained a F1-score of 88.29%, improving over the second-best model by about 1.9%. Its advantage was even clearer on the clinically heterogeneous TUAR corpus, where it reached a F1-score of 85.70% and outperformed the closest competitor by approximately 3.0%. On the more balanced AAR dataset, G3DCT performed on par with the best baseline model, with nearly identical F1 scores (90.46% vs. 90.52%), while achieving the highest recall (90.73%). We further analyzed performance on combined artifacts, which constitute 20.38% of MotionArt and 14.74% of TUAR (Table 3). On MotionArt, G3DCT achieved the highest F1-score of 87.43%, outperforming the second-best baseline model by about 1.9%. On TUAR, G3DCT again attained the best F1-score of 78.98%, surpassing RNN-CNN by approximately 2.2%. To gain further insight into the decision process of G3DCT, we visualized the scalp attention maps produced by the Fast Scalp Attention Module (FSAM). As illustrated in Fig. 5, regions with high attention weights align closely with the known physiological origins of specific artifact types, indicating that the model learns to focus on physiologically meaningful spatial patterns.

**Table 1.** Artifact detection performance on the MotionArt dataset.

| Model | Acc | Pre | Rec | F1 | Spe | BAC |
| --- | --- | --- | --- | --- | --- | --- |
| G3DCT | **0.9730** | **0.9978** | **0.9621** | **0.9797** | **0.9956** | **0.9789** |
| CLAttn | 0.9581 | 0.9906 | 0.9469 | 0.9683 | 0.9814 | 0.9641 |
| TCN | 0.9150 | 0.9711 | 0.9010 | 0.9348 | 0.9441 | 0.9226 |
| GAT | 0.9230 | 0.9806 | 0.9040 | 0.9408 | 0.9627 | 0.9334 |
| RNN-CNN | 0.9609 | 0.9922 | 0.9496 | 0.9705 | 0.9845 | 0.9670 |
| LDA | 0.8550 | 0.9489 | 0.8301 | 0.8855 | 0.9068 | 0.8684 |

**Table 2.** Multi-class artifact classification performance across datasets.

| Model | MotionArt |  |  |  |  | TUAR |  |  |  |  | AAR |  |  |  |  |
| --- | --- | --- | --- | --- | --- | --- | --- | --- | --- | --- | --- | --- | --- | --- | --- |
|  | ACC | Pre | Rec | F1 | W-F1 | ACC | Pre | Rec | F1 | W-F1 | ACC | Pre | Rec | F1 | W-F1 |
| G3DCT | **0.9278** | **0.8676** | **0.9055** | **0.8829** | **0.9287** | **0.9021** | **0.8311** | **0.8901** | **0.8570** | **0.9044** | 0.9042 | 0.9046 | **0.9073** | 0.9046 | 0.9046 |
| CLAttn | 0.8721 | 0.7867 | 0.8237 | 0.7987 | 0.8750 | 0.8910 | 0.7897 | 0.8769 | 0.8267 | 0.8942 | 0.8098 | 0.8345 | 0.8190 | 0.8184 | 0.8154 |
| TCN | 0.8585 | 0.7686 | 0.8293 | 0.7904 | 0.8653 | 0.8693 | 0.7359 | 0.8826 | 0.7862 | 0.8768 | 0.8279 | 0.8427 | 0.8399 | 0.8320 | 0.8278 |
| GAT | 0.5554 | 0.4590 | 0.4615 | 0.4094 | 0.5801 | 0.6053 | 0.4348 | 0.6529 | 0.4531 | 0.6549 | 0.4693 | 0.5652 | 0.4758 | 0.4558 | 0.4517 |
| RNN-CNN | 0.9169 | 0.8621 | 0.8773 | 0.8660 | 0.9172 | 0.8912 | 0.8264 | 0.8446 | 0.8346 | 0.8917 | **0.9048** | **0.9053** | 0.9070 | **0.9052** | **0.9056** |
| LDA | 0.9073 | 0.8479 | 0.8242 | 0.8350 | 0.9062 | 0.6028 | 0.3538 | 0.3136 | 0.3118 | 0.5866 | 0.8062 | 0.8145 | 0.8111 | 0.8110 | 0.8091 |

**Table 3.**
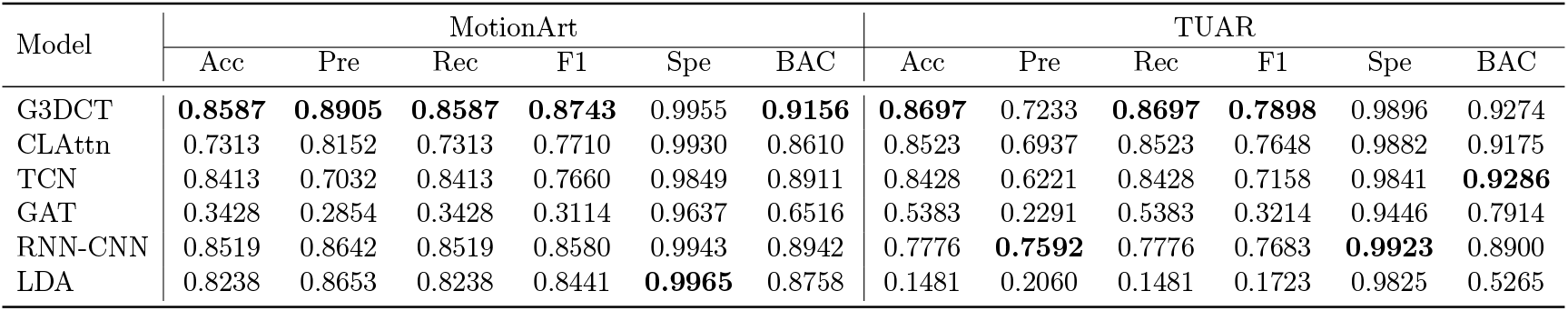
Classification performance on combined-artifact categories.

**Figure 5.**
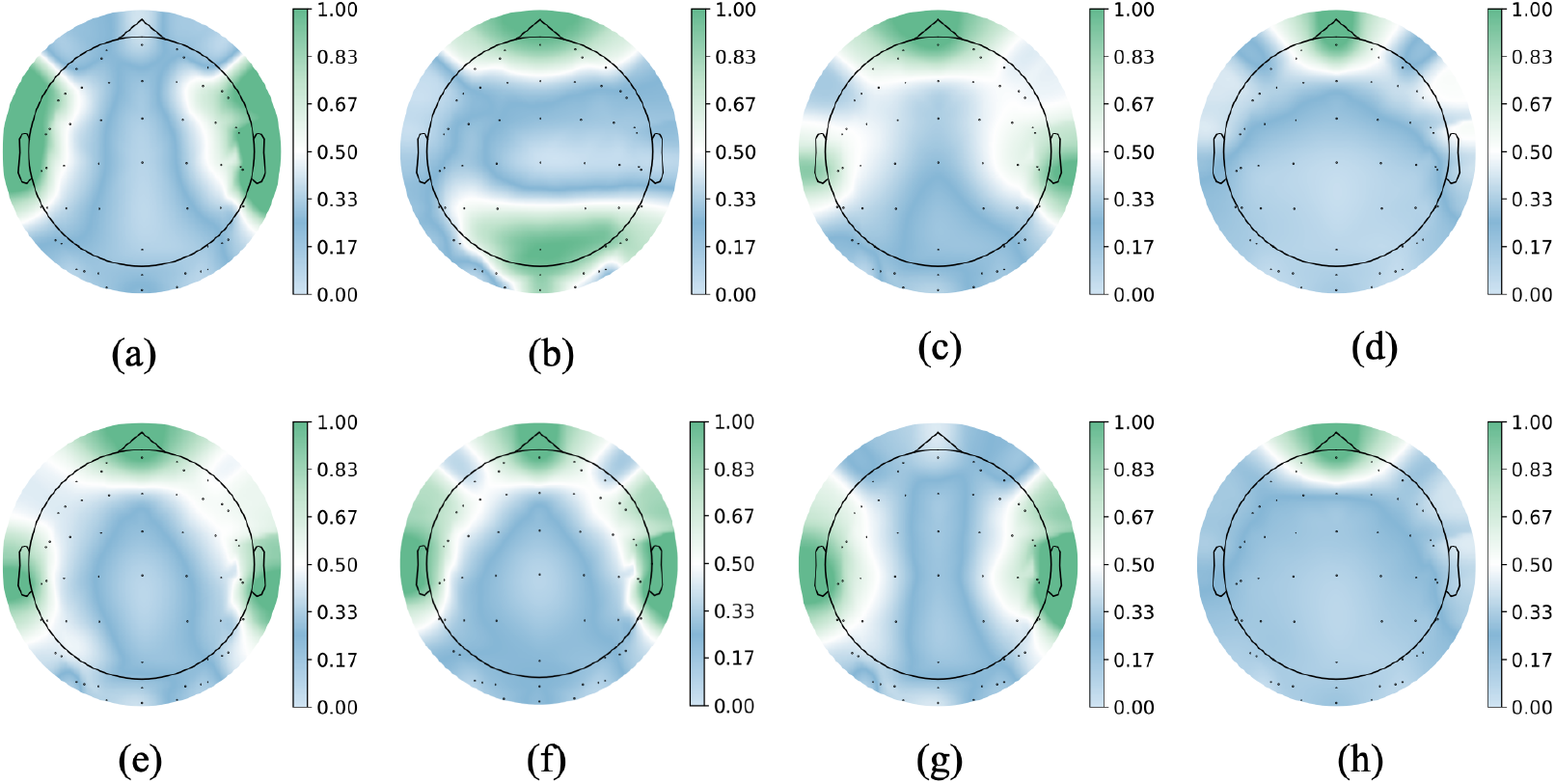
Spatial attention patterns learned by the FSAM module. (a) Chewing; (b) Vertical head movement; (c) Vertical eye movement; (d) Blinking and eyebrow movement; (e) Tongue movement and eyebrow movement; (f) Swallowing and eyebrow movement; (g) Swallowing; (h) Blinking.

### 4.2. Sensitivity Analysis

To assess the robustness of the optimal hyperparameter configuration 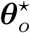 identified via TPE, each key hyperparameter was varied individually around its optimal value. For each setting, performance was evaluated through multiple independent training runs across different outer folds and random seeds. The results are summarized in Fig. 6. Consistent with prior studies 52, 53, excessively small learning rates (e.g., 10^−5^) substantially slowed convergence and reduced final accuracy, whereas overly large values (e.g., 5 *×* 10^−3^) frequently caused training instability. In contrast, variations in the focal-loss *γ*, dropout rate, and *α*-scale factor within the tested ranges did not lead to significant performance degradation. These results indicate that while the learning rate requires careful tuning, G3DCT remains robust to moderate variations in the other hyperparameters.

**Figure 6.**
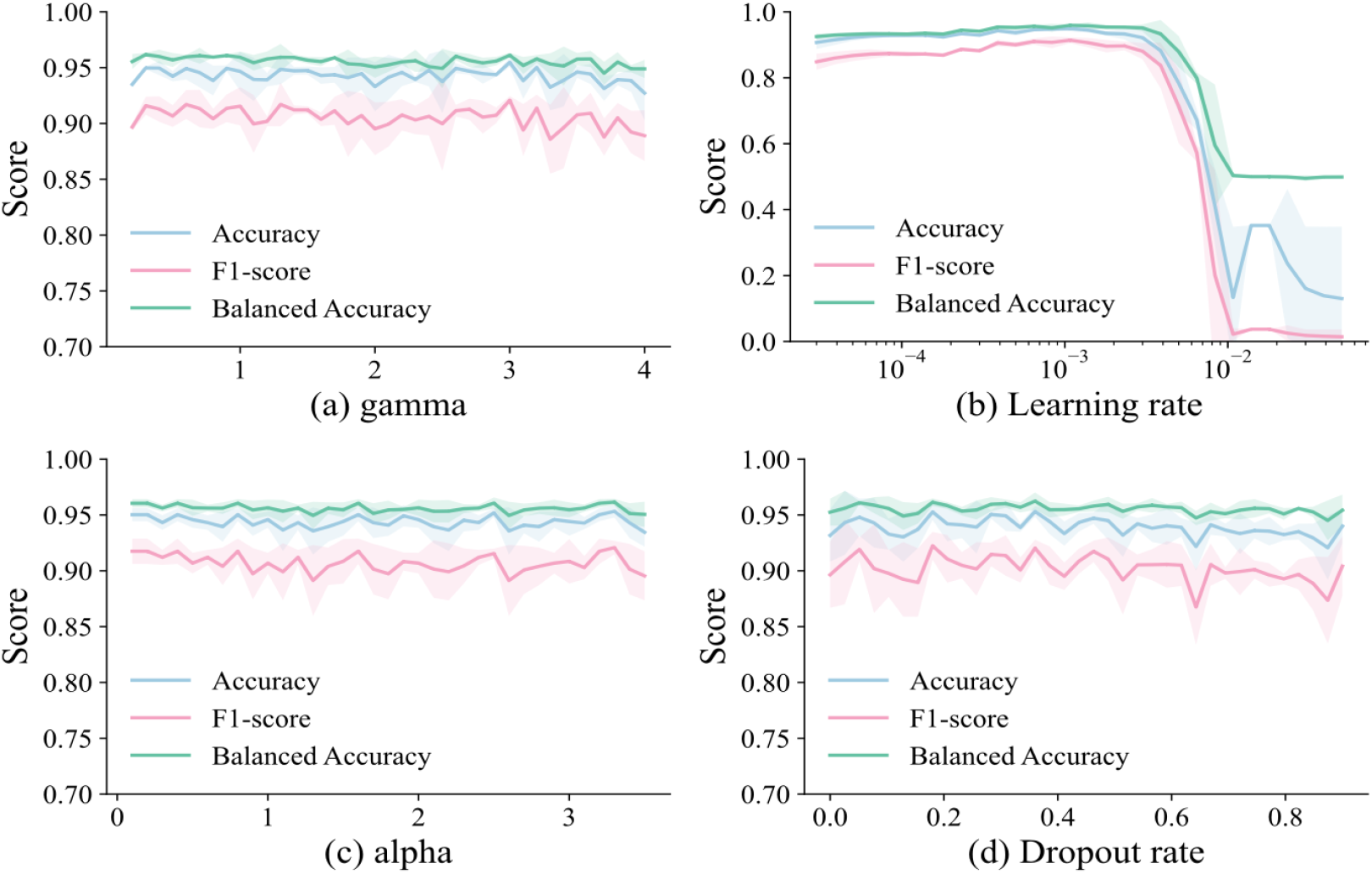
Hyperparameter sensitivity analysis of G3DCT. Curves report the mean performance across folds, and the shaded areas indicate the 95% confidence interval.

### 4.3. Ablation Studies

To clarify the contribution of each component in the proposed G3DCT architecture, we conducted a systematic ablation study on the multi-class classification task using the MotionArt dataset. The overall results are summarized in Table 4, where the full model (A0) is evaluated under the best-performing hyperparameter configuration obtained from the tuning procedure. Module-level ablations indicate that each core component contributes positively to the final performance. Removing the 3D-CNN blocks (A1) caused the most pronounced degradation among architectural modules (a 13.3% drop in F1-score), highlighting the importance of local spatiotemporal feature extraction when the electrode topology is explicitly preserved in the input representation. Disabling the Temporal Convolutional Pyramid (TCP) module (A2) also decreased the F1-score to 0.8553 (a 9.2% drop), confirming its effectiveness in modeling multi-scale temporal dynamics. Similarly, removing the Transformer encoder (A3) impaired performance, resulting in an F1-score of 0.9076 (a 3.7%reduction), suggesting that attention-based temporal modeling offers complementary benefits beyond convolutional temporal encoding. Input-structure ablations validate the necessity of preserving electrode topology. Flattening the input into a 2D channel–time representation (B1) reduced the F1-score to 0.8549 (a 9.3% drop), while randomly shuffling the channel order (B2) led to a comparable decline to 0.8528 (a 9.5% drop). These results demonstrate that explicitly encoding spatial topology is a key factor that improves the discriminative capability. Finally, training-strategy ablations highlight the impact of loss design and optimization choices. Removing class weighting (C1) reduced the F1-score to 0.8692 (a 7.7% drop), indicating that training which accounts for class distribution is necessary for stable multi-class recognition. Both fixed class weights (C2, F1 = 0.8980) and standard cross-entropy loss (C3, F1 = 0.9004) underperformed compared with the proposed configuration, reflecting the advantage of the adaptive weighting strategy for handling imbalanced distributions. Replacing the optimizer with Adam (C4) further degraded performance (an 8.6% drop), suggesting that the optimization setting also impacts convergence quality.

**Table 4.** Ablation study results.

| Experiment | Acc | Pre | Rec | F1 | Spe | BAC |
| --- | --- | --- | --- | --- | --- | --- |
| A0 | **0.9606** | **0.9411** | **0.9451** | **0.9421** | **0.9969** | **0.9710** |
| A1 | 0.8883 | 0.8165 | 0.8300 | 0.8170 | 0.9913 | 0.8300 |
| A2 | 0.9063 | 0.8596 | 0.8630 | 0.8553 | 0.9926 | 0.9278 |
| A3 | 0.9418 | 0.9019 | 0.9215 | 0.9076 | 0.9955 | 0.9215 |
| B1 | 0.9144 | 0.8498 | 0.8721 | 0.8549 | 0.9933 | 0.9327 |
| B2 | 0.9106 | 0.8564 | 0.8634 | 0.8528 | 0.9930 | 0.9282 |
| C1 | 0.9303 | 0.8924 | 0.8596 | 0.8692 | 0.9945 | 0.9270 |
| C2 | 0.9367 | 0.9043 | 0.9061 | 0.8980 | 0.9951 | 0.9506 |
| C3 | 0.9411 | 0.9103 | 0.9007 | 0.9004 | 0.9953 | 0.9007 |
| C4 | 0.9110 | 0.8599 | 0.8774 | 0.8615 | 0.9931 | 0.8774 |

A0: full model; A1: removing 3D-CNN; A2: removing TCP; A3: removing Transformer; B1: 2D flattened input; B2: with channel order shuffled; C1: without class weights; C2: with fixed class weights; C3: using cross-entropy loss; C4: using Adam optimizer.

## 5. Discussion

Through systematic experiments and comparative evaluations, this study demonstrates that joint spatiotemporal modeling constitutes an effective strategy for the robust decoding of EEG signals 54. The proposed Grid-based 3D Convolution-Transformer (G3DCT) framework achieved superior perfor-mance across three public datasets, offering an optimal balance among identification accuracy, computational efficiency, and physiological interpretability. Specifically, the framework offers the following advantages. First, it introduces a method to overcome device variability by mapping each electrode to a unified 9 *×* 9 grid template. This unified representation enables the model to maintain stable performance while adapting to other devices. Second, while maintaining comparable or even better classification performance, this lightweight architecture significantly reduces inference resource consumption. Compared to the multi-branch ensemble model (RNN-CNN), it uses only 21% of the parameters and model size, achieves an inference throughput of approximately 19.3k samples per second, reduces peak GPU memory usage from about 2.0 GB to about 0.8 GB, and nearly halves the per-fold training time. Third, the model leverages Fast Scalp Attention Module (FSAM) in combination with the electrode topology, generating heatmaps that align closely with physiological structure. Specifically, blink and vertical eye movement artifacts primarily activate the prefrontal regions 55. Horizontal eye movement artifacts show symmetric distribution over the left and right frontotemporal areas 56. Chewing artifacts are concentrated in channels over the temporalis muscles 57. Such physiologically consistent heatmaps enhance model interpretability, facilitating its potential use in clinical practice. Despite these advantages, the study revealed that the model exhibits confusion in distinguishing between single and combined artifacts. For instance, when presented with an artifact involving both tongue movement and eyebrow raising, the model tends to misclassify it solely as eyebrow movement. We posit that this confusion arises when one component of the combined artifacts (e.g., frontal EMG) exhibits an excessively strong signal 58. In long-sequence analyses, this dominant signal may overshadow weaker features (e.g., from tongue activity).

Comparative analysis further confirms that effective spatiotemporal joint modeling relies on a complementary combination of local spatial features and global temporal modeling 59, 60. Although RNN-CNN achieves high accuracy by integrating these two strengths, this benefit comes at the cost of parallel execution of multiple branches during inference. This results in increased latency and GPU memory consumption as the number of branches grows, thereby limiting its flexibility for real-world deployment 61. In comparison, static graph methods (e.g., GAT) exhibit limitations in both spatial and temporal dimensions. Spatially, they depend on a predefined adjacency graph to model relationships between electrodes 48. However, in practical applications, scenarios such as channel missing or addition can render the predefined graph structure ineffective or inapplicable 62. In terms of temporal modeling, the node features in such methods typically rely on hand-crafted statistical features and temporal signals processed by shallow networks 63. This simplified temporal representation has limited capacity, making it difficult to capture complex, long-range temporal dependencies in the signals. On the MotionArt dataset, the combination of handcrafted features and a linear classifier achieves competitive results. However, its recognition performance declines significantly on the other two datasets, which validates the earlier concern about the limited cross-dataset robustness of handcrafted feature-based models.

The CLAttn and TCN models demonstrate competent performance in temporal modeling, yet their capabilities in spatial modeling remain limited. These approaches often treat signals from each electrode as independent time series, failing to adequately exploit the spatial relationships between electrodes7, 64. Consequently, these models struggle to distinguish between artifacts with similar spectral signatures. A prime example is differentiating eyebrow raising (a frontal artifact) from chewing (which affects temporal areas), both being mid-to-high frequency myogenic artifacts58. In terms of temporal modeling, TCN that relies on increasing the dilation rate to expand its receptive field can lead to the loss of local details40. Meanwhile, LSTMs often suffer from state decay or update overwriting in long sequences, resulting in inadequate retention of long-range contextual information65, 66. The G3DCT framework is designed to directly address these limitations. To overcome the shortcomings in spatial modeling, we employ a grid mapping strategy based on the physical electrode layout, combined with 3D convolution that simultaneously extracts features across channel, spatial neighborhood, and temporal dimensions67. This approach effectively leverages the distinct spatial distribution patterns of different artifacts. For temporal modeling, we introduce a hybrid architecture integrating a temporal convolutional pyramid with pre-normalized Transformers, which successfully mitigates the inherent limitations of both TCN and LSTM41.

In summary, this study systematically demonstrates the critical role of integrating spatial structure, multi-scale temporal information, and long-range dependencies for EEG artifact recognition. We demonstrate that embedding spatial information into the model’s internal structure effectively mitigates the impact of variations in acquisition devices and referencing schemes. On the practical side, the unified grid mapping and lightweight model proposed in this work enhance deployment flexibility. The heatmaps generated by FSAM exhibit a clear correspondence with brain anatomical regions, enhancing the interpretability in real-world applications. In brain-computer interface and real-time intervention scenarios, the low inference latency and more stable recognition capability for minor classes contribute to building more reliable systems.

In addition to the previously mentioned issues, the model has other limitations. While the 9 *×* 9 grid serves as a spatial normalization method to align data from different devices, it may also lead to information loss 68. For systems with too few channels or unique layouts, spatial details may be oversimplified. For high-density EEG setups (e.g., 128 or 256 channels), the grid may struggle to fully preserve spatial patterns 69. Furthermore, the fixed-length window segmentation approach offers limited effectiveness in modeling slow artifacts that last significantly longer than the window length 70. In response to the above limitations, following directions can be focussed on in future work. The 9 *×* 9 grid can be enhanced by introducing variable-resolution grids (adaptively selecting higher or lower resolutions based on channel count) and learnable coordinate projection (trainable mapping from 2D electrode coordinates to the grid), thereby better accommodating electrode layouts of varying densities to improve spatial modelling 68. For combined artifact recognition, multi-label learning can be explored to enable the model to identify multiple coexisting artifacts in the signal 71. Regarding long-term modeling and segmentation strategies, artifact recognition paradigms based on object detection and tracking can be investigated 70, 72. This approach will no longer rely on predefined time segments but will instead directly localize the onset, duration, and offset of artifacts in continuous EEG signals, enabling dynamic capture and full characterization of long-duration artifacts. These efforts are directed toward evolving this framework into an interpretable and practical artifact management tool, with future validation planned through large-scale studies in clinical and home environments 73.

## 6. Conclusion

This study introduced Grid-based 3D Convolution-Transformer (G3DCT), an interpretable framework for EEG artifact identification that jointly models the spatial arrangement of electrodes and temporal signal dynamics. By standardizing EEG recordings into a 9 *×* 9 spatial grid, G3DCT achieves consistent learning of spatial relationships across different montages and acquisition devices. Based on this representation, a lightweight hybrid architecture effectively captures multi-scale local features as well as long-range temporal dependencies, proving especially advantageous for detecting challenging combined artifacts. Furthermore, the proposed Fast Spatial Attention Module (FSAM) generates physiologically meaningful scalp attention maps, providing an intuitive basis for model interpretation and facilitating practical clinical verification.

Comprehensive evaluations on three public datasets demonstrate that G3DCT achieves strong overall performance while maintaining favorable computational efficiency. These results indicate that G3DCT holds practical promise as a robust and deployable tool for automated EEG quality control. Potential real-world applications include clinical monitoring, brain–computer interface systems, and long-term wearable EEG recording.

## CRediT authorship contribution statement

**Aonan He**: Writing – original draft, Visualization, Investigation, Data curation, Methodology, Conceptualization. **Xi Wang**: Writing – review & editing, Methodology. **Jiangwei Yu**: Investigation, Data curation. **Xiaojia Wang**: Writing – review & editing, Supervision. **Zongyuan Ge**: Writing review & editing, Supervision, Conceptualization. **Youyong Kong**: Resources, Writing – review & editing, Supervision. **Guanyu Yang**: Writing review & editing, Supervision. **Chunfeng Yang**: Project administration, Writing – review & editing, Investigation, Conceptualization, Funding acquisition. **Chen Yang**: Project administration, Writing – review & editing. **Miao Cao**: Project administration, Writing – original draft, Investigation, Conceptualization, Funding acquisition.

## Declaration of competing interest

The authors declare that they have no known competing financial interests or personal relationships that could have appeared to influence the work reported in this paper.

## Acknowledgments

This work was supported by the National Key Research and Development Program of China [grant number 2023YFC3603600] and the National Natural Science Foundation of China [grant numbers T2225025, 31400842]. The authors thank the Big Data Computing Center of Southeast University for providing the facility on the numerical calculations in this paper. MC acknowledges the facilities and scientific and technical assistance of the Australian National Imaging Facility, a National Collaborative Research Infrastructure Strategy (NCRIS) capability, at the Swinburne Neuroimaging Facility, Swinburne University of Technology. MC acknowledges research fellowship fundded by Australian National Health and Medical Research Council (NHMRC) Ideas Grant [Grant number 2038089].

## Data availability

The TUAR and AAR datasets analyzed in this study are publicly available from their official sources: the [TUH EEG (NEDC) repository](https://isip.piconepress.com/projects/nedc/html/tuh_eeg/) and the [AAR dataset repository](https://github.com/stefan-ehrlich/dataset-automaticArtifactRemoval). The MotionArt dataset collected by the authors is publicly available on the OpenNeuro (ds006386). The source code and trained model checkpoints are available at GitHub (AonanH/G3DCT).

## Notes

### Competing Interest Statement

The authors have declared no competing interest.

