## Supplementary Material for "G3DCT: An Interpretable Spatial Grid-based Framework with Temporal Convolution-Transformer for EEG Artifact Identification"

#### S1 Dataset channel configurations

To clarify the variability of EEG channel configurations across datasets, we summarize the electrode montages used in MotionArt, TUAR, and AAR. MotionArt was collected using a relatively dense montage with **59 EEG channels**, covering frontal, central, temporal, parietal, and occipital regions (Table S1).

Table S1: Channel configuration of the MotionArt dataset (59 channels).

| **MotionArt channels (total = 59)** | | | |
|---|---|---|---|
| Fpz | Fp1 | Fp2 | AF3 |
| AF4 | AF7 | AF8 | Fz |
| F1 | F2 | F3 | F4 |
| F5 | F6 | F7 | F8 |
| FCz | FC1 | FC2 | FC3 |
| FC4 | FC5 | FC6 | FT7 |
| FT8 | Cz | C1 | C2 |
| C3 | C4 | C5 | C6 |
| T7 | T8 | CP1 | CP2 |
| CP3 | CP4 | CP5 | CP6 |
| TP7 | TP8 | Pz | P3 |
| P4 | P5 | P6 | P7 |
| P8 | POz | PO3 | PO4 |
| PO5 | PO6 | PO7 | PO8 |
| Oz | O1 | O2 | |

In contrast, TUAR exhibits substantial heterogeneity in acquisition devices and recording protocols. Different subsets contain different channel sets, and only **19 channels** are consistently shared across the three representative folders (`01_tcp_ar`, `02_tcp_le`, `03_tcp_ar_a`) (Table S2).

Table S2: Common EEG channels across different TUAR subsets.

| **TUAR subset** | **Common channels** |
|---|---|
| `01_tcp_ar` (23) | A1, A2, C3, C4, CZ, F3, F4, F7, F8, FP1, FP2, FZ, O1, O2, P3, P4, PZ, T1, T2, T3, T4, T5, T6 |
| `02_tcp_le` (27) | 30, A1, A2, C3, C4, CZ, EKG, F3, F4, F7, F8, FP1, FP2, FZ, O1, O2, OZ, P3, P4, PG1, PG2, PZ, T3, T4, T5, T6, PHOTIC PH |
| `03_tcp_ar_a` (23) | 29, 30, 31, 32, C3, C4, CZ, F3, F4, F7, F8, FP1, FP2, FZ, O1, O2, P3, P4, PZ, T3, T4, T5, T6 |
| **Intersection of all three subsets** (19) | C3, C4, CZ, F3, F4, F7, F8, FP1, FP2, FZ, O1, O2, P3, P4, PZ, T3, T4, T5, T6 |

The AAR dataset has a fixed montage after removing EOG channels, resulting in a consistent **27-channel** configuration for all recordings (Table S3).

Table S3: Channel configuration of the AAR dataset after removing EOG channels (27 channels).

| **AAR channels (total = 27)** | | |
|---|---|---|
| Fp1 | Fz | F3 |
| F7 | FC5 | FC1 |
| C3 | T7 | CP5 |
| CP1 | Pz | P3 |
| P7 | O1 | O2 |
| P4 | P8 | CP6 |
| CP2 | Cz | C4 |
| T8 | FC6 | FC2 |
| F4 | F8 | Fp2 |

### S2 Label distribution

We further summarize the label distributions of the three datasets to highlight the class imbalance.

Table S4: Label distribution of the MotionArt dataset (total $N = 16637$).

| **Label** | **Count** | **Proportion (%)** |
|---|---|---|
| blink | 5916 | 35.6 |
| chew | 1640 | 9.9 |
| eyebrow | 1474 | 8.9 |
| blink_hor_headm | 1438 | 8.6 |
| hor_headm | 1253 | 7.5 |
| tongue | 1025 | 6.2 |
| ver_headm | 698 | 4.2 |
| blink_ver_headm | 695 | 4.2 |
| blink_eyebrow | 625 | 3.8 |
| ver_eyem | 537 | 3.2 |
| tongue_eyebrow | 515 | 3.1 |
| hor_eyem | 502 | 3.0 |
| swallow | 202 | 1.2 |
| swallow_eyebrow | 117 | 0.7 |

Table S5: Label distribution of the TUAR dataset (total $N = 107919$).

| **Label** | **Count** | **Proportion (%)** |
|---|---|---|
| musc | 41924 | 38.8 |
| eyem | 25909 | 24.0 |
| elec | 22563 | 20.9 |
| musc_elec | 7781 | 7.2 |
| eyem_musc | 6582 | 6.1 |
| chew | 1617 | 1.5 |
| chew_elec | 570 | 0.5 |
| eyem_elec | 553 | 0.5 |
| eyem_chew | 420 | 0.4 |

Table S6: Label distribution of the AAR dataset ( total $N = 6631$).

| **Label** | **Description** | **Count** | **Proportion (%)** |
|---|---|---|---|
| S4 | Smooth horizontal eye movement | 910 | 13.7 |
| S6 | Smooth vertical eye movement | 910 | 13.7 |
| S10 | Head flexing | 780 | 11.8 |
| S7 | Talking | 780 | 11.8 |
| S5 | Saccadic vertical eye movement | 651 | 9.8 |
| S2 | Eyes closed | 650 | 9.8 |
| S3 | Saccadic horizontal eye movement | 650 | 9.8 |
| S8 | Blinking | 650 | 9.8 |
| S9 | Clenching | 650 | 9.8 |

### S3 Implementation Details of Baseline Methods

#### S3.1 CNN–LSTM with Attention

CLAttn is a lightweight temporal modeling baseline that combines a 1D convolutional front-end with a recurrent sequence encoder and an attention-based aggregation mechanism. Given an input EEG segment $\mathbf{X} \in \mathbb{R}^{T \times C}$ (time length $T$ and channel number $C$), the model first applies a two-layer 1D CNN along the temporal dimension to extract local temporal patterns from multi-channel signals. The extracted feature sequence is then fed into a multi-layer LSTM to capture longer-range temporal dependencies. Finally, a learnable attention module assigns importance weights to each time step and produces a weighted sum representation, which is passed to a fully connected classifier for artifact label prediction.

#### S3.2 Temporal Convolutional Network

The Temporal Convolutional Network (TCN) is a purely convolutional baseline designed for sequence modeling. Given an EEG segment $\mathbf{X} \in \mathbb{R}^{T \times C}$, the model applies a stack of causal 1D convolutional residual blocks with exponentially increasing dilation factors, enabling a large temporal receptive field while preserving the temporal order. Each temporal block consists of two dilated Conv1D layers followed by nonlinear activation and dropout, together with a residual connection for stable optimization. The final sequence representation is obtained by global average pooling over the temporal dimension, and a linear classification head outputs the predicted artifact label.

#### S3.3 Graph Attention Network

The Graph Attention Network (GAT) is a graph-based baseline that models EEG electrodes as nodes and their neighborhood relations as edges. In our implementation, we constructed a fixed graph topology using a 4-neighborhood adjacency defined on a $9 \times 9$ electrode grid. For each EEG segment, node features were computed from the raw signals by extracting simple channel-wise statistics (mean and standard deviation), resulting in a feature vector for each electrode. The model applies multiple stacked GAT convolution layers to propagate and aggregate information across connected electrodes via attention-weighted message passing. A global mean pooling layer is then used to obtain a graph-level representation, followed by a multilayer perceptron classifier to predict the artifact class.

#### S3.4 RNN–CNN Ensemble

The RNN–CNN baseline is an ensemble model that combines recurrent and convolutional temporal feature extractors for EEG artifact classification. Given an EEG segment represented as a multivariate time series ($T \times C$), the RNN branch adopts an LSTM to capture long-range temporal dependencies across time steps, and the final hidden state is fed into a fully-connected classifier to produce class logits. In parallel, two independent 1D-CNN branches are constructed to learn hierarchical temporal patterns via stacked convolution, normalization, activation, and max-pooling layers, followed by fully-connected classification layers. The final prediction is obtained by averaging the logits from the three branches (one LSTM-based model and two CNN-based models), forming a simple yet effective late-fusion ensemble.

#### S3.5 LDA with Hand-crafted Features

The LDA baseline is a classical machine-learning approach that performs EEG artifact classification based on hand-crafted statistical and spectral features. For each EEG segment, we extract a fixed-length feature vector from every channel, including time-domain statistics (e.g., mean, standard deviation, variance, skewness, kurtosis, RMS, and zero-crossing rate), frequency-domain band power features (delta/theta/alpha/beta/gamma), and additional complexity measures (e.g., spectral entropy, Hjorth parameters, and approximate entropy). All channel-wise features are concatenated to form a global representation of the input segment. Before classification, the features are optionally reduced using univariate feature selection (ANOVA F-test), and then normalized by standardization. Finally, a Linear Discriminant Analysis (LDA) classifier is trained to predict artifact classes in the resulting feature space.

### S4 Hyperparameter Search Space and Best Configuration

Table S7: Hyperparameter search space and best configuration on the MotionArt dataset.

| Hyperparameter | Search Range / Candidates | Best Configuration |
|---|---|---|
| Learning rate ($lr$) | loguniform $[5 \times 10^{-5}, 3 \times 10^{-3}]$ | $9.6569 \times 10^{-4}$ |
| Weight decay | loguniform $[1 \times 10^{-6}, 3 \times 10^{-4}]$ | $7.4054 \times 10^{-5}$ |
| Batch size | $\{64, 128, 256\}$ | 128 |
| Transformer layers | $\{2, 3\}$ | 2 |
| Transformer heads | $\{4, 8\}$ | 4 |
| FFN dimension | $\{256, 512, 768\}$ | 512 |
| Reduce dimension | $\{128, 256\}$ (divisible by #heads) | 128 |
| FC dimension | $\{256, 384\}$ | 384 |
| Dropout | uniform $[0.2, 0.5]$ | 0.3384 |
| Epochs / Patience | epochs = 20, patience = 5 | epochs = 20, patience = 5 |
| Focal $\gamma$ | uniform $[1.0, 2.5]$ | 2.3598 |
| $\alpha$ scale | uniform $[0.7, 1.6]$ | 0.8329 |
| Per-class scaling | $\forall c:$ uniform $[0.6, 1.4]$ | [1.1381, 1.3799, 1.2600, 0.7958, 1.1164, 1.1524, 1.0874, 0.6579, 1.2859, 1.2800, 1.0600, 1.2309, 0.6235, 0.7800] |

Table S8: Table S8: Hyperparameter search space and best configuration on the TUAR dataset.

| Hyperparameter | Search Range / Candidates | Best Configuration |
|---|---|---|
| Learning rate ($lr$) | loguniform $[1 \times 10^{-6}, 5 \times 10^{-4}]$ | $3.7751 \times 10^{-4}$ |
| Weight decay | loguniform $[1 \times 10^{-6}, 5 \times 10^{-4}]$ | $1.2407 \times 10^{-6}$ |
| Batch size | $\{128, 256\}$ | 256 |
| Transformer layers | $\{2, 3\}$ | 2 |
| Transformer heads | $\{4, 8\}$ | 8 |
| FFN dimension | $\{256, 512, 768\}$ | 768 |
| Reduce dimension | $\{128, 256\}$ (divisible by #heads) | 128 |
| FC dimension | $\{256, 384\}$ | 384 |
| Dropout | uniform $[0.2, 0.4]$ | 0.3715 |
| Epochs / Patience | epochs = 40, patience = 8 | epochs = 40, patience = 8 |
| Focal $\gamma$ | uniform $[1.0, 2.0]$ | 1.9512 |
| $\alpha$ scale | uniform $[0.9, 0.999]$ | 0.9913 |
| Per-class scaling | minor classes ($c \in \mathcal{C}_{\text{minor}}$): uniform $[1.0, 1.5]$; mid-frequency classes ($c \in \mathcal{C}_{\text{mid}}$): uniform $[0.9, 1.2]$; major classes ($c \in \mathcal{C}_{\text{major}}$): uniform $[0.8, 1.1]$ | [0.8423, 1.0438, 1.1335, 1.2690, 1.0613, 1.1938, 1.3028, 1.2545, 1.1731] |

Table S9: Hyperparameter search space and best configuration on the AAR dataset.

| Hyperparameter | Search Range / Candidates | Best Configuration |
|---|---|---|
| Learning rate ($lr$) | loguniform $[5 \times 10^{-5}, 3 \times 10^{-3}]$ | $1.9697 \times 10^{-4}$ |
| Weight decay | loguniform $[1 \times 10^{-6}, 3 \times 10^{-4}]$ | $1.4726 \times 10^{-4}$ |
| Batch size | $\{64, 128, 256\}$ | 64 |
| Transformer layers | $\{2, 3\}$ | 3 |
| Transformer heads | $\{4, 8\}$ | 4 |
| FFN dimension | $\{256, 512, 768\}$ | 512 |
| Reduce dimension | $\{128, 256\}$ (divisible by #heads) | 128 |
| FC dimension | $\{256, 384\}$ | 384 |
| Dropout | uniform $[0.2, 0.5]$ | 0.3846 |
| Epochs / Patience | epochs = 20, patience = 5 | epochs = 20, patience = 5 |
| Focal $\gamma$ | uniform $[1.5, 2.5]$ | 2.3573 |
| $\alpha$ scale | uniform $[0.99, 0.999]$ | 0.9950 |
| Per-class scaling | $\forall c:$ uniform $[0.9, 1.99]$ | [1.7147, 1.5638, 1.8958, 0.9006, 0.9642, 1.8688, 1.6090, 1.0725, 1.9164] |

### S5 Confusion Matrices Across Datasets and Baseline Models

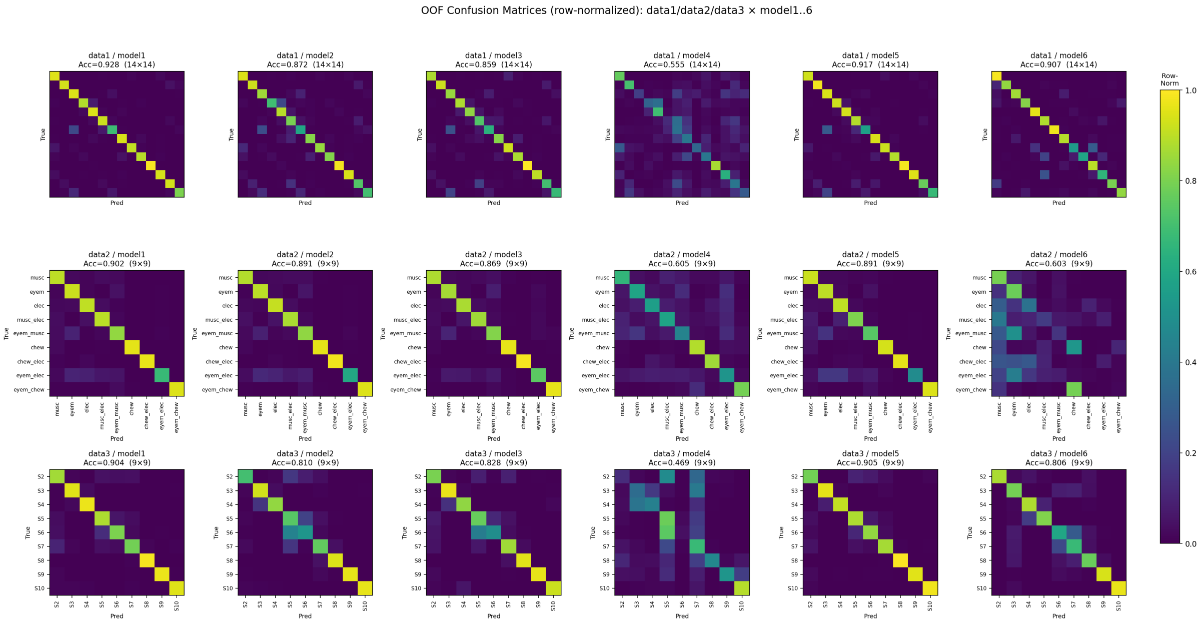


Figure S1: Datasets are arranged by rows: MotionArt (data1, 14 classes), TUAR (data2, 9 classes), and AAR (data3, 9 classes). Models are arranged by columns: G3DCT (model1), CLAttn (model2), TCN (model3), GAT (model4), RNN–CNN ensemble (model5), and LDA (model6).
